# Variable Resolution Maps (VRM) in CCTBX and Phenix: Accounting For Local Resolution In cryoEM

**DOI:** 10.64898/2026.03.25.714315

**Authors:** Pavel V. Afonine, Paul D. Adams, Alexandre G. Urzhumtsev

## Abstract

Calculation of density maps from atomic models is essential for structural studies using crystallography and electron cryo-microscopy (cryoEM). These maps serve various purposes, including atomic model building, refinement, visualization, and validation. However, accurately comparing model-calculated maps to experimental data poses challenges, particularly because the resolution of cryoEM experimental maps varies across the map. Traditional crystallography methods generate finite-resolution maps with uniform resolution throughout the unit cell volume, while most modern software in cryoEM employ Gaussian-like functions to generate these maps, which does not adequately account for atomic model parameters and resolution. Recent work by Urzhumtsev & Lunin (2022, *IUCr Journal*, 9, 728-734) introduces a novel method for computing atomic model maps that incorporate local resolution and can be expressed as analytically differentiable functions of all atomic parameters. This approach enhances the accuracy of matching atomic models to experimental maps. In this paper, we detail the implementation of this method in CCTBX and Phenix.

**Synopsis:** New tools implemented in CCTBX and Phenix allow the calculation of variable-resolution maps through a sum of atomic images expressed as analytic functions of all atomic parameters, along with their associated local resolution.

## 1. Introduction

Calculation of density maps from atomic models is fundamental to structural studies in crystallography and electron cryo-microscopy (cryoEM). These maps represent the distribution of electron density (X-ray crystallography), nuclear density (neutron crystallography), or electrostatic scattering potential (electron diffraction or cryoEM) for a given model. In structural biology, model-calculated maps are used for various goals and some recent examples are atomic model building (Mostosi *et al*., 2020; Pfab *et al*., 2021), flexible fitting (Trabuco *et al*., 2008; Kirmizialtin *et al*., 2015), refinement (DiMaio *et al*., 2009, 2015; Afonine *et al*., 2018b), map improvement (He *et al*., 2023), ligand identification (Robertson *et al*., 2020), validation (Afonine *et al*., 2018a; Pintilie *et al*., 2020), structure prediction (Si *et al*., 2020), and computer molecular graphics (Meng *et al*., 2023). In this context, model-calculated maps are either matched against experimentally derived maps to assess, or enhance the fit of the atomic model to observed data, or serve as training data for deep learning applications, or as temporary objects to speedup calculations (Sayre, 1951). Consequently, the ability to accurately and efficiently compute density maps from atomic models is important.

A proper comparison between a model and experimental data requires that both are expressed in compatible terms. Since experimental maps always have finite resolution, model-calculated maps must match this resolution. In crystallography, finite-resolution maps are usually computed by a three-steps procedure. First, the exact density distribution is generated by summing individual atomic densities. Then corresponding Fourier coefficients are calculated and truncated at the experimental resolution limit. Finally, this truncated set of Fourier coefficients is converted back into a map that exhibits a uniform resolution aligned with that of the experimental data. This procedure is hardly applicable in cryoEM where the resolution of the experimental map is usually varies over the map (e.g., Cardone et al., 2013).

Modern software typically generates atomic model maps by summing contributions from individual atoms within a specified surrounding volume. Most programs use Gaussian-like functions (Mooij *et al*., 2006; Topf *et al*., 2008; DiMaio *et al*., 2009, 2015; Kirmizialtin *et al*., 2015; Pintilie *et al*., 2020; Zhang *et al*., 2020; Blau *et al*., 2023; He *et al*., 2023; Meng *et al*., 2023; Cai *et al*., 2025; Selvaraj *et al*., 2025), following the work of Costain (1941) and Booth (1946), who noted that “*Examination of the shape of the atomic peaks derived from a number of Fourier syntheses shows that the radial density distribution can be closely represented by the [Gaussian] function*.” Additionally, machine learning techniques can also be employed to generate model maps (Zhang *et al*., 2025). These model map calculations present two issues. First, they do not explicitly account for atomic model parameters such as occupancy, atomic displacement parameters (ADPs), chemical element types, and charges. DiMaio *et al*. (2015) utilize Agarwal’s (1978) expression for atomic electron density while disregarding atomic occupancy. Second, these models assume that atomic shapes at finite resolution manifest solely as positive density peaks, aligning with Diamond’s (1971) assertion that “*Theoretically, therefore, the image of an isolated atom … will not be distinguishable from Gaussian to an extent worth characterizing*.” In reality, finite resolution atomic images oscillate with both positive and negative ripples. This becomes evident when the experimental map is displayed with both positive and negative contours, in contrast to the customary positive-only representation. The central peak corresponding to the Fe ion in the cryo-EM map for mouse heavy-chain apoferritin (Kucukoglu *et al*., 2024), along with the subsequent dip, clearly illustrates this point (Fig. 1a, compare blue and orange lines in Fig. 1b). Failing to account for these ripples in the model map can degrade molecular images (Fig. 1c in Urzhumtsev & Lunin, 2022) and result in incorrect values of the atomic displacement parameter obtained by real-space refinement of atomic models (Lunin & Lunina, 2025). To address this problem, Chapman (1995, 2013) employed atomic images derived from finite-resolution Fourier transforms of piecewise approximations to the respective atomic scattering functions, though this method appears computationally intensive.

**Figure 1.**
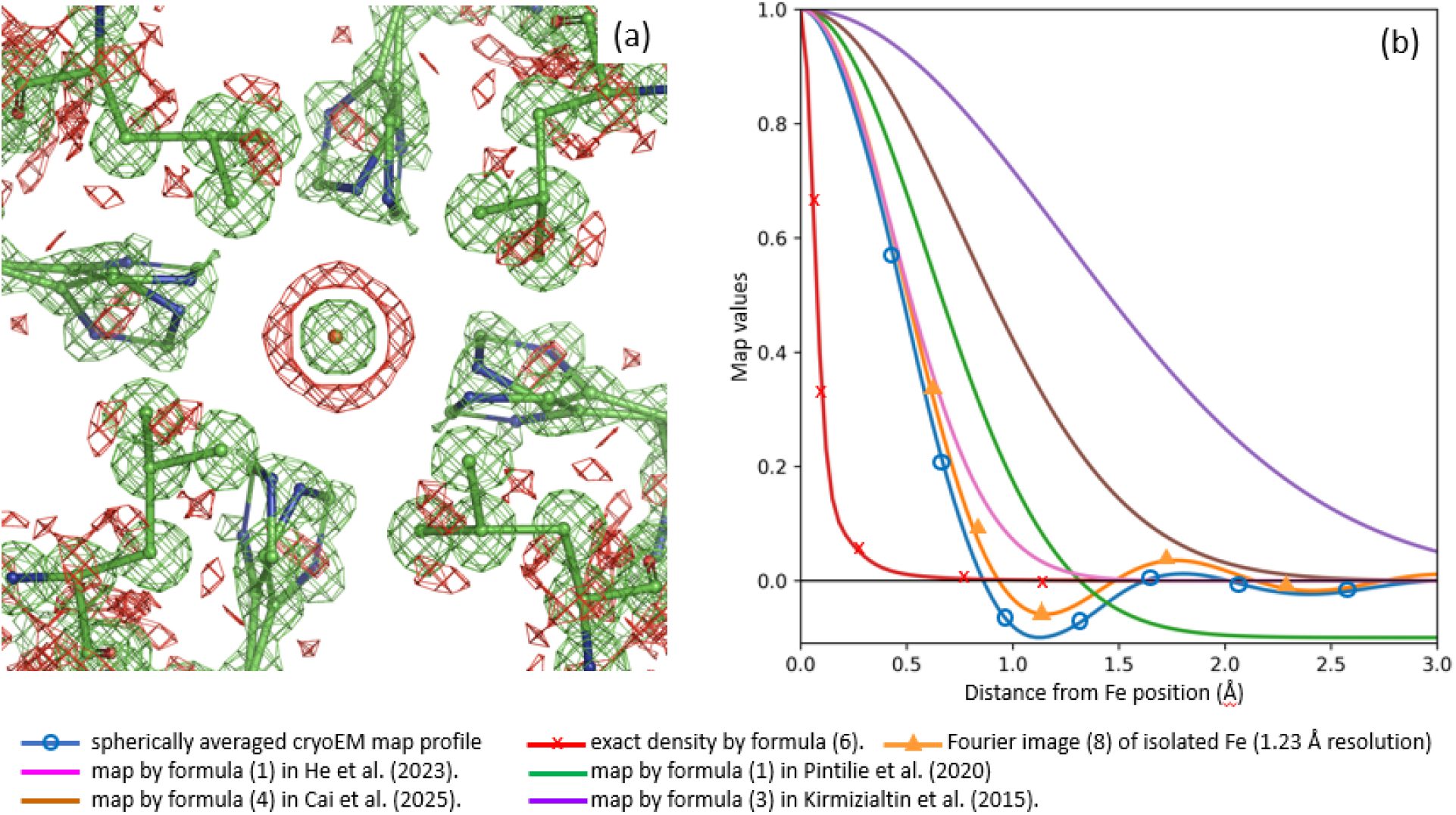
(a) CryoEM model of mouse heavy-chain apoferritin (PDB: 8RQB), focusing on Fe 201 in chain A, and corresponding experimental map (EMDB: 19436) contoured at ±1.5 units. (b) Radial components of the experimental cryoEM map around Fe (blue curve), exact electron density (red curve) and its finite-resolution image (orange curve), and of several maps calculated for an isolated Fe atom. All maps normalized by their values at the origin (maximum value).

Recently, Urzhumtsev & Lunin (2022) introduced a method that allows for the computation of atomic model maps that account for local resolution and are expressed as analytically differentiable functions of all atomic parameters. The latter readily enables their use in real-space refinement and machine learning training. In this approach, the entire variable-resolution model map is computed as a sum of individual atomic contributions, each one representing the atom’s image at a specified resolution approximated by

Here the function

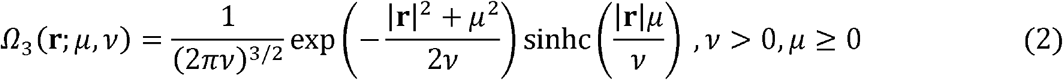

(Urzhumtsev & Lunin, 2024) can describe both the peak in the origin and Fourier ripples,**r**_*n*_ is the center, *q*_*n*_ is the occupancy, and *B*_*n*_ is the isotropic displacement parameter of the atom *n*. For a given resolution *d*_*min*_, coefficients *A*^(*k*)^, *B*^(*k*)^, and *R*^(*k*)^ depend solely on the atomic type and can be pre-computed and tabulated.

In this work, we describe the implementation of the proposed method in CCTBX (Grosse-Kunstleve *et al*., 2002) and Phenix (Liebschner *et al*., 2019). This implementation paves the way for its use in applications requiring finite-resolution atomic model-generated maps and makes the method accessible to the broader community.

## 2. Method

### 2.1. Images of atomic density

Originating from Vand *et al*. (1957), multi-Gaussian approximations to the atomic scattering functions are widely used, in its current form proposed by Agarwal (1978)

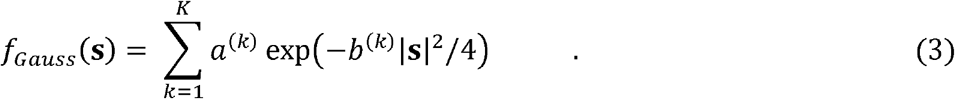

Here **s** is a scattering vector, *s* = |**s**| = 2 sin *θ*/*λ* is its norm, and coefficients *a*^(*k*)^ and *b*^(*k*)^ are specific to the chemical element and the experiment type.

To model isotropic structural disorder, atomic densities of immobile atoms are convoluted with the Gaussian functions

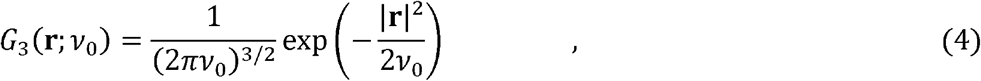

where *v*_ο_ is related to the isotropic atomic displacement parameter *B* as *v*_ο_ = *B*/8*π*^2^. Due to the convolution property of Gaussians

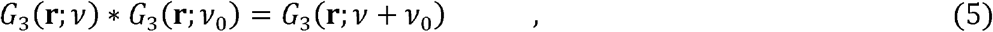

respective approximations to the atomic densities calculated with (3) get a form (Agarwal, 1978)

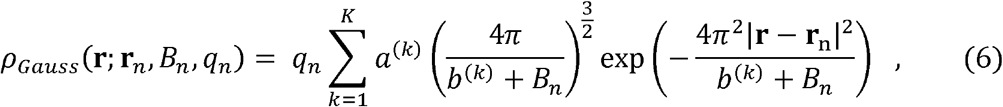

conventionally used to generate atomic density by their sums. Due to spherical symmetry of all functions considered here, in what follows, we deal with their radial components which we note by overline, *e*.*g*.,

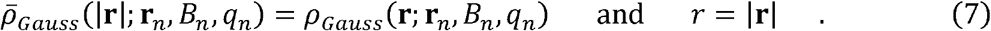

The radial component of the finite resolution Fourier image of the density (6) for an atom with a unit occupancy and placed in the origin is calculated as

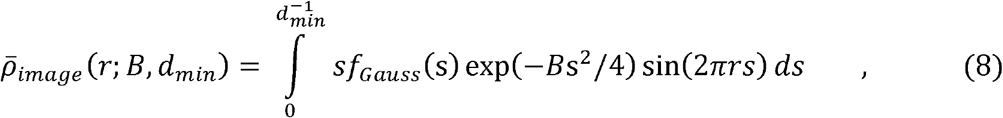

where *d*_*min*_ is the resolution cutoff, except for *r* ≪ 1 where it is calculated as

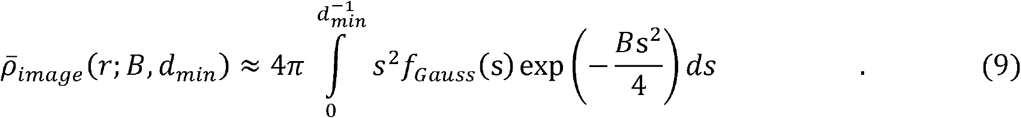

These images were used as the reference for obtaining *A*^(*k*)^, *B*^(*k*)^, and *R*^(*k*)^ in (1).

### 2.2. Normalizing atomic images

Images for different atoms at different resolutions look drastically different. Their magnitude is proportional to the number of electrons (for electron densities), and the positions of extrema of the radial components spaced apart approximately at *d*_*min*_ distance. Consequently, *B*^(*k*)^ values grow as a square of a distance between these peaks. While at resolutions around 1 Å these coefficients are of the order of ten, at lower resolutions, *B*^(*k*)^ are two to three orders of magnitude larger (Fig. 2a). Generally, numerical optimization favors variables being on the same scale, while may behave poorly when it is not the case.

**Figure 2.**
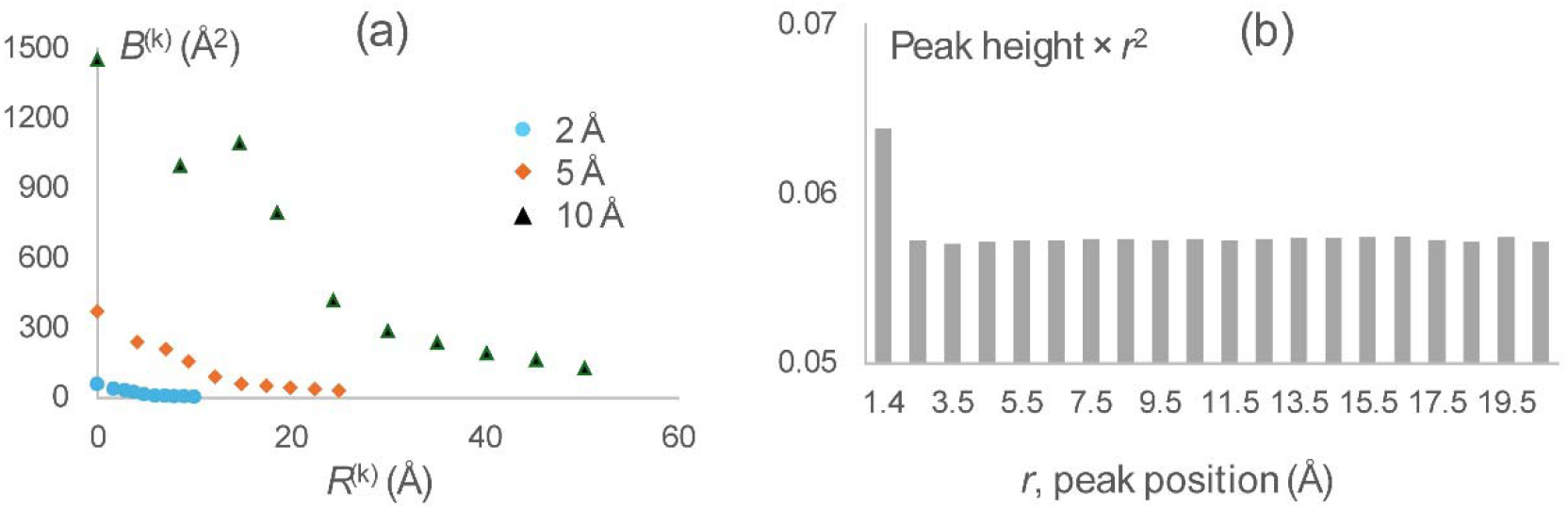
(a) Ten first coefficients *R*^(*k*)^ and *B*^(*k*)^ of the approximation to the images of a carbon atom at the resolution 2, 5, and 10 Å. (b) Height of the ripples in the 1 Å – resolution image of a carbon atom normalized by the value in its origin. The height is shown as a function of the distance to the origin, being multiplied by the square of this distance.

We find that the finite resolution images of immobile atoms expressed as a function of a normalized, dimensionless parameter *q* = *r*/*d*_*min*_ appear similar to that of *d*_*min*_ = 1Å resolution. Moreover, an additional scaling of the curve magnitude to its value at the origin as

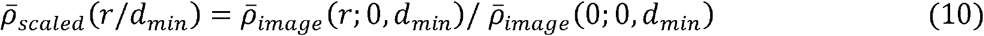

makes the normalized atomic images (10) similar to each other for all atomic types and for all resolutions (Fig. 3).

**Figure 3.**
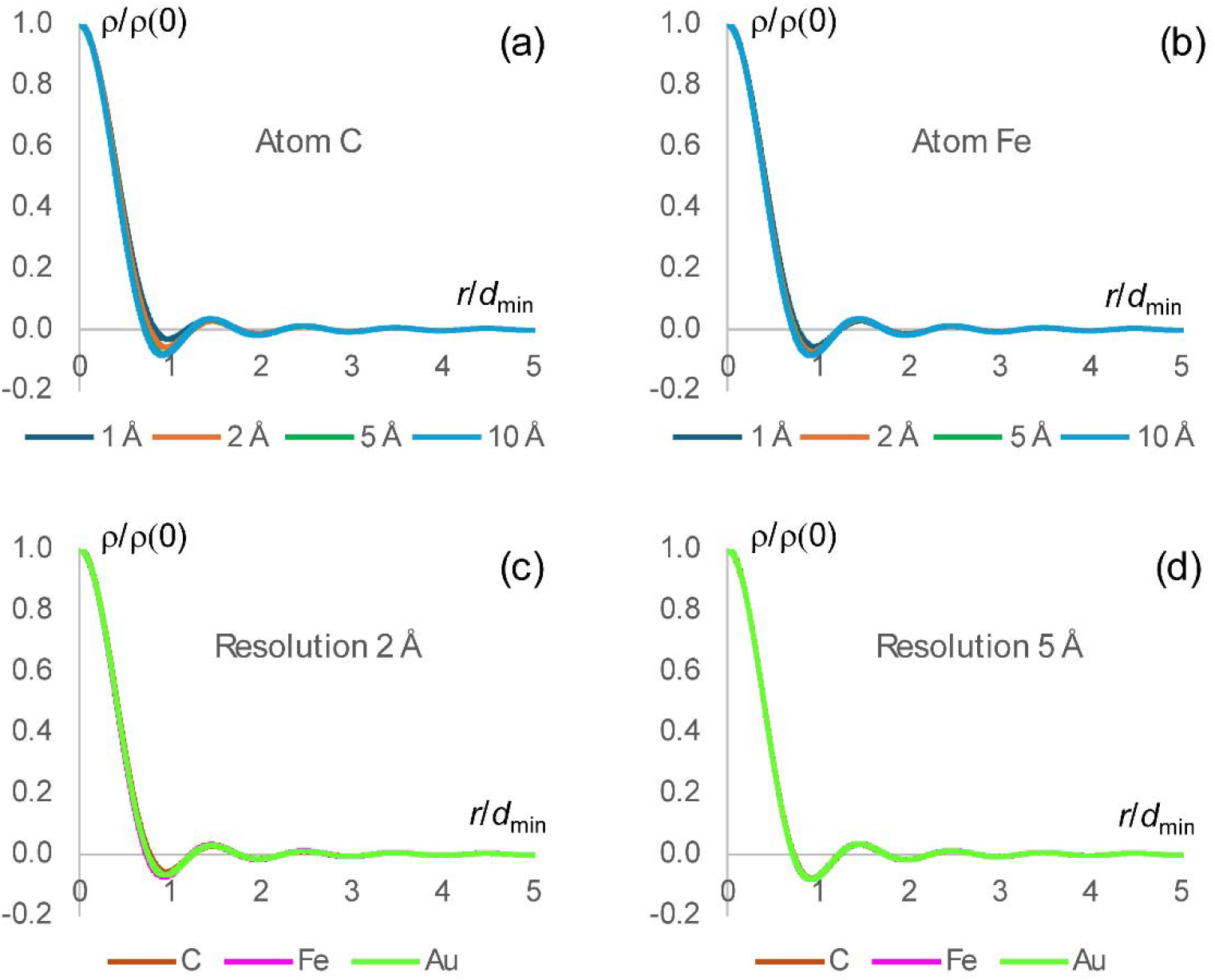
Atomic images scaled as in (10). Panels (a) and (b) show images for carbon (C) and iron (Fe) atoms at different resolutions. Panels (c) and (d) display images for carbon, iron, and gold (Au) atoms at resolutions of 2 Å and 5 Å, respectively. Notably, despite the significant differences in both the types of atoms and the resolutions used in this example, all curves for the scaled images virtually coincide.

The utility of such image scaling is two-fold. First, expectedly, it drastically improves the numerical procedures of the search for coefficients of the approximation (1) to the rescaled image (10). When the approximation coefficients 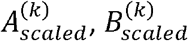, and 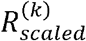 for a normalized curve 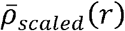 (10) at the interval [0;*q*_*max*_]are determined, the coefficients for the original curve (1), that is on the absolute distance scale, in Å, and for the distance up to *r*_*max*_ = *q*_*max*_*d*_*min*_, are obtained by reverting the scaling:

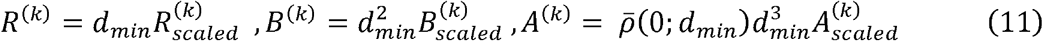

(see, *e*.*g*., equation (6) to understand dependence of coefficients *A*^(*k*)^ and *B*^(*k*)^ on the distance scale). To keep the total integral of the function (“atomic charge”), the *q*_*max*_ *= r*_*max*_/*d*_*min*_ value has been recommended to be integer or half-integer (Urzhumtsev *et al*., 2022).

Second, the similarity of the scaled images allows reusing their corresponding *A*^(*k*)^, *B*^(*k*)^, and *R*^(*k*)^ coefficients, once found for a given atoms at given resolution, as a starting point for the search of coefficients for different atoms or/and resolutions, which greatly improves the results.

### 2.3. Approximation to finite resolution atomic images

In this work, finite-resolution maps are calculated following (1) where *Ω*_3_(**r**; *μ, ν*) (2) appears instead of Gaussians in (6). In fact, Gaussian function (4) is a particular case of the function *Ω*_3_(**r**; *μ, ν*) corresponding to *μ* = 0. This function has its shape concentrated in a shell around a sphere of radius *μ* with a width defined by the parameter *ν*. The key feature of these spherical shell functions is that, similar to (5), they do not change their shape after convolution

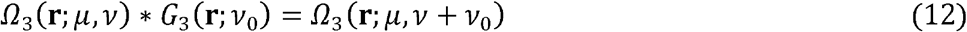

with the Gaussian function (Urzhumtsev & Lunin, 2024). This is especially important for modeling atomic images with positional uncertainties, as this is done for atomic densities. Based on the properties of *Ω*_3_(**r**; *μ, ν*), two approaches to calculate an approximation (1) to atomic images have been proposed (Urzhumtsev *et al*., 2022).

In the first approach, for an atom of a given type with *B*_*n*_ = 0 (immobile atom) and a unit occupancy, one calculates its image *ρ*_*n*_ (**r**;*d*_*min*_)placed at the origin at the required resolution *d*_*min*_, and then builds an approximation (1) to this image as

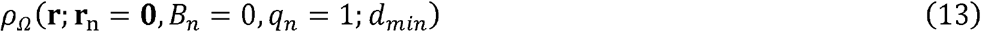

Approximation (1) to an atomic image at the same resolution with any *B*_*n*_ ≠ 0 is then available using (1), (2) and (10). By construction, the constants *A*^(*k*)^,*B*^(*k*)^, and *R*^(*k*)^ are the same for all atoms of the same chemical type.

In the second approach, one builds the *Ω*_3_(**r**; *μ, ν*) approximation (1) to the three-dimensional unit interference function

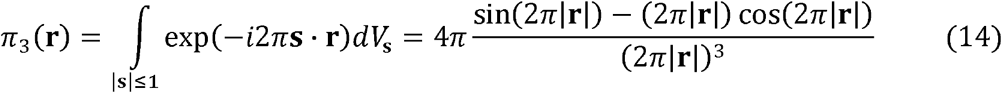

which is the image of a point scatterer at the resolution *d*_*min*_ = 1 Å (Urzhumtsev & Lunin, 2024). With (10) and other properties of *Ω*_3_(**r**; *μ, ν*) and of the interference function, this provides the coefficients of the approximation (1) to the image of any Gaussian function (4) at any chosen resolution *d*_*min*_ (Urzhumtsev & Lunin, 2024). Combining these images according to the *K*-Gaussians approximation (6) to the atomic densities, this gives the coefficients of an approximation (1) to the image of any atom with any isotropic *B*_*n*_ value at any resolution *d*_*min*_ as a sum of the respective Gaussian images.

This latter approach, implemented in *ModQMap* program (Urzhumtsev & Lunin, 2025), is more general, because it computes a unique set of coefficients analytically adjustable to any resolution, while is less attractive computationally. The number of terms for an image of each individual Gaussian component is roughly the same as the number of coefficients for the whole atomic image in the former approach. Thus, the number of terms in (1) is about *K*-times more (usually about 5-6 times) compared with such number for the former approach; the same for the CPU time. Since CPU time is especially important for refinement where the calculation of maps and its derivatives is repeated many times, we programmed the former method of the approximation.

### 2.4. Approximation protocol

#### 2.4.1. Obtaining approximation parameters for a given image

The set of parameters *A*^(*k*)^,*B*^(*k*)^, and *R*^(*k*)^ for an image of a given atom at the specified resolution *d*_*min*_ are obtained by fitting function (1) to the reference atomic image (8–9) as described by Urzhumtseva *et al*. (2023). This process is governed by two principal parameters: *q*_*max*_, defining the distance limit *r*_*max*_ = *q*_*max*_*d*_*min*_ from the atom’s center, up to which the approximation is calculated, and Δ*ρ*_*max*_, the maximum allowed approximation error, that is computed as the absolute value of the difference between the reference curve and its approximation calculated over the interval [0,*r*_*max*_]. Their values, and the coefficients *a*^(*k*)^ and *b*^(*k*)^ of the scattering function (3) are the input parameters to the procedure.

Calculation of optimal *A*^(*k*)^,*B*^(*k*)^, and *R*^(*k*)^ occurs in two steps: assignment of approximate values of the coefficients, according to the curve’s extrema position and shape, followed by their refinement to minimize the least-squares discrepancy between the reference atomic image (8-9) and its approximation (1). The scaled atomic image 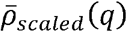,*q* ≥ 0, at a given resolution *d*_*min*_ is analyzed in increasing order of the argument value from 0 up to the distance limit slightly beyond *q*_*max*_ (see Section 2.4.3); then, for a given position of a local extremum, a new term is noted if Δ*ρ*_*max*_ is above the predefined maximal allowed deviation (see also Section 2.4.3). After the whole interval [0,*q*_*max*_] is analyzed, the curve calculated using (1) is subtracted from the reference curve. Optionally, more iterations can be done when the residual curve is scanned for local extrema again, and if any found above Δ*ρ*_*max*_ additional term is added that optimally fits it.

Numeric minimization of the least-squares difference between the reference curve and its approximation (1) is done by L-BFGS-B method (Liu & Nocedal, 1989), resulting in the coefficients 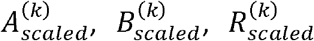 for the scaled image. As it is typical for a local optimization, the accuracy of the approximation with these coefficients largely depends on their initial values. This creates a room for several combinations of initial assignment of the coefficient values and their following refinement to find the best approximation. There are three algorithms used to initiate the procedure:

1. parameters of all assigned terms are refined after the whole curve being analyzed;
2. parameters of all terms are refined after the whole curve being analyzed, except the parameters of the first term approximating the central peak which are refined as soon as they are estimated;
3. parameters of each term are refined, together with the previously defined parameters, as soon as they are estimated.

The computation time in each of the three algorithms is dominated by that of refinement of the coefficient values, increasing with the number of the coefficients. By this reason, the third algorithm requires more CPU time while sometime gives a more accurate approximation.

At the end of the procedure, the coefficients *A*^(*k*)^,*B*^(*k*)^, *R*^(*k*)^ for the original, unscaled atomic image are calculated according to (11).

#### 2.4.2. Tabulating parameter values

For a given atom, coefficients of the approximation can either be calculated as needed for a specific resolution *d*_*min*_ or pre-calculated and tabulated in advance for a set of atoms (*e*.*g*., the entire periodic table) and for a range of resolution values, allowing them to be reused later as necessary. The latter may be useful for highly repetitive calculations such as atomic model refinement, where on-the-fly calculation of image approximations may quickly become a computational bottleneck. For tabulating over the resolution range from 1 up to 2 Å, the step 0.005 Å has been chosen such that when moving between adjacent resolutions, the value of atomic image in the origin changes by less than 1% of the peak value. Beyond 2 Å the step can be kept at 0.01 Å which is the accuracy of the typically reported resolution values.

Calculating approximation coefficients for a series of images (*e*.*g*., when tabulating them) has an additional advantage due to the similarity of the scaled images (10), which can help in finding an approximation more accurately and faster compared to determining the approximation for a single atom at a single resolution. Approximate values of 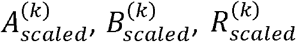 for a scaled image of a given atom at a given resolution may be taken as the initial values for refinement for the image of the same atom calculated at the adjacent resolution or as those for the image of another atom at the same resolution.

Figure 4 outlines the procedure of finding optimal *A*^(*k*)^,*B*^(*k*)^, and *R*^(*k*)^ for a given atom at a series of resolutions. The procedure starts from the determination of the set of coefficients for the starting resolution value, as described in Section 2.4.1, assuming the resolution range is ordered by value from smallest to largest. Then it cycles over subsequent resolution values in the range. For each new resolution, the refined coefficients for the scaled image at a previous resolution are used as the starting values for refinement. This drastically accelerates refinement since usually these values are closer to the final values than the starting values defined by algorithms 1 – 3 (Section 2.4.1). At the same time, it is necessary to check if such repeated recycling of initially defined coefficient values does not miss a better set of coefficients. For this, we additionally reuse, the algorithms 1 – 3 (faster algorithms 1 and 2 are applied more frequently) and select the best result among all tested options.

**Figure 4.**
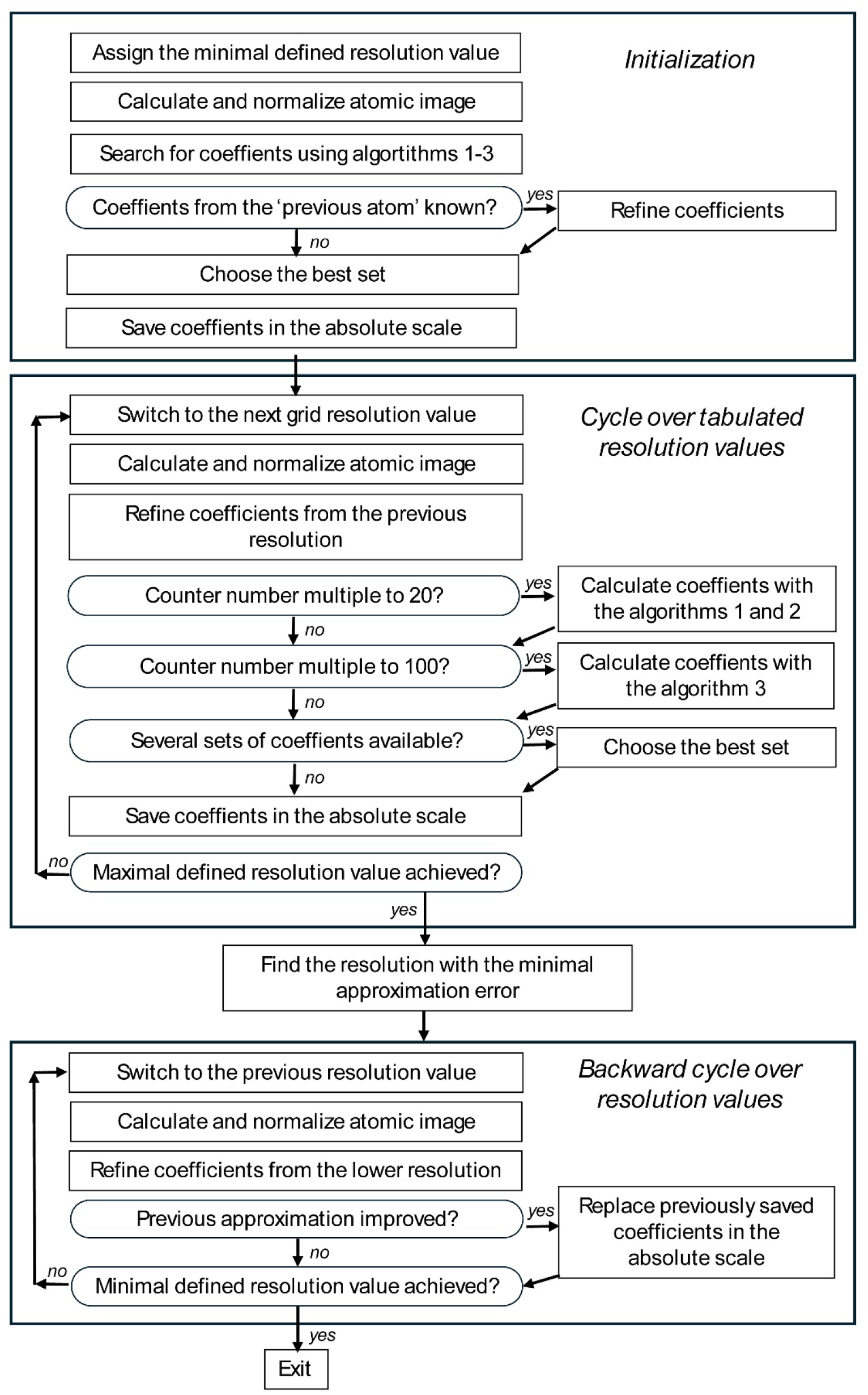
Flowchart of the procedure to determine the approximation coefficients (1) for a set of finite-resolution images of an atom of a given chemical type.

In this protocol the coefficients for higher resolutions pass through a much smaller number of refinement iterations that those for lower resolution and therefore may be less accurate. To correct this, we scan over the whole set of resolutions for which the approximation coefficients have been calculated and identify the set of coefficients giving the minimal approximation error. Then, starting from this resolution and corresponding coefficients, the procedure is running in the opposite sense, taking the coefficients from the lower resolution and refining them to the higher resolution. If a better result is obtained, which is usually the case, the coefficients found during the forward resolution analysis are replaced by these new values.

As this is done moving from one resolution to another, a similar approach is applicable moving from one atomic type to another. Optionally, and complementary to algorithms 1 – 3, the initialization step of the procedure for each atomic type may also try a set of refined coefficients 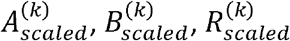 from an atom processed previously.

#### 2.4.3. Number of approximation terms

The number of terms in approximation (1) represents a trade-off between the runtime cost of the map calculation and the desired accuracy of the approximation. While more terms improve accuracy, fewer terms lead to faster calculations. This is especially important since atomic images decrease with distance much slower than the densities themselves (Fig. 2b), thus necessitating a larger distance cut-off value *r*_*max*_ than it is required to compute the exact density (Urzhumtsev *et al*., 2022), and, consequently, a larger number of grid nodes for which the calculations should be done.

Intuitively, at least one term in (1) is required to represent the peak at the origin and at least one term is necessary to model each Fourier ripple, with two ripples per each distance interval of the length of approximately *d*_*min*_. For such an approximation, the total number of terms with *R*^(*k*)^≤ r_*max*_ for an image at the resolution of *d*_*min*_ is about *K* = 2··(*r*_*max*_/*d*_*min*_) = 2*q*_*max*_. In fact, the atomic images (8) are calculated and analyzed up to the distance *r*_*extended*_ that is slightly larger than *r*_*max*_ = *q*_*max*_*d*_*min*_. The extended approximation interval is used because it may happen that including one or a few terms with *R*^(*k*)^ >*r*_*max*_ may notably reduce approximation errors at the end of the interval [0;*r*_*max*_]. For example, for the image of a carbon atom at the resolution *d*_*min*_ = 2Å, working with the approximations up to the distance limit *r*_*max*_ = 5Å (*q*_*max*_ = 2.5), the two terms around are *R*^(5)^ = 4.86 and *R*^(6)^ = 5.95 (Table 1). The series cut at *R*^(5)^ Fig. 5a, black curve) gives much larger errors at the upper boundary of the interval [0;*r*_*max*_]than a series completed by *R*^(6)^ = 5.95 (Fig. 5a, red curve). Arrows of the respective color indicate these *R*^(*k*)^ values.

**Table 1.**
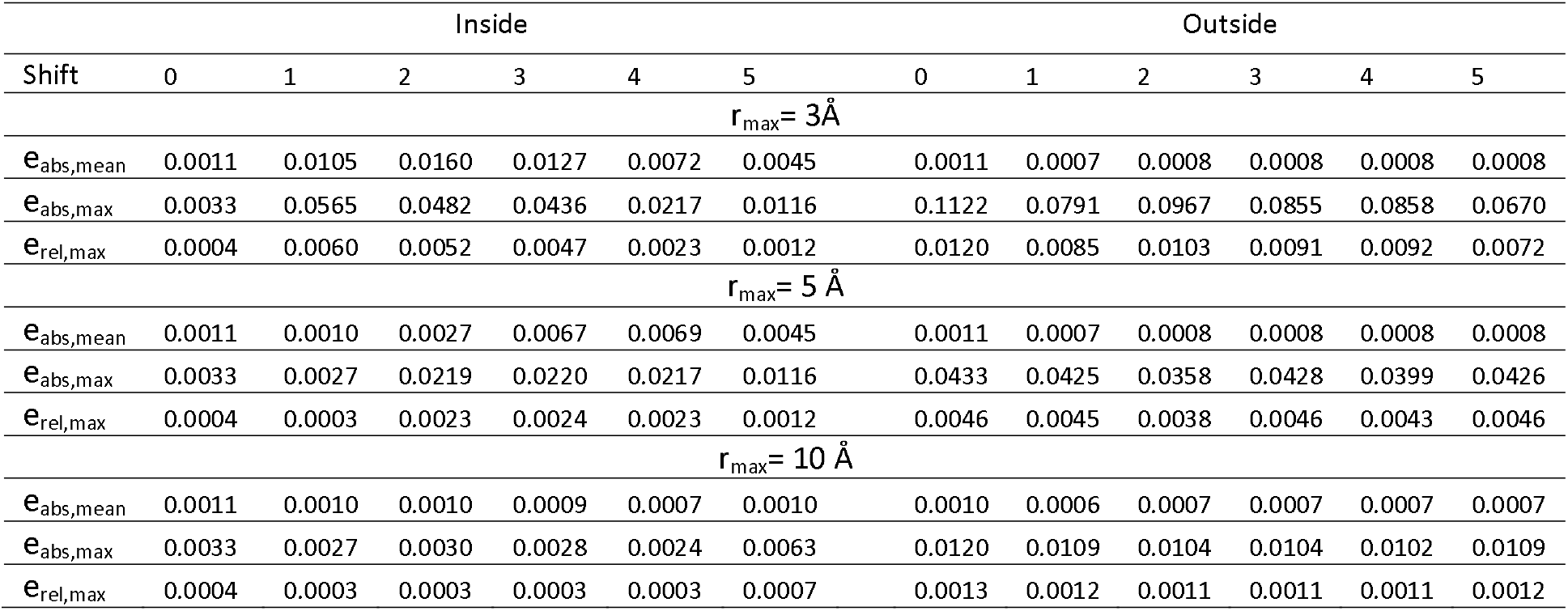
Discrepancy measures between *ρ*_*FT*_ and *ρ*_*image*_ maps for six two-atom models. Statistics shown separately for the points inside and outside the molecular region. Error metrics are defined in §3.1. Shift defines the distance between the two atoms by the respective number of Ångstroms following each of the three axes (see §3.2.1).

|  | Inside |  |  |  |  |  | Outside |  |  |  |  |  |
| --- | --- | --- | --- | --- | --- | --- | --- | --- | --- | --- | --- | --- |
| Shift | 0 | 1 | 2 | 3 | 4 | 5 | 0 | 1 | 2 | 3 | 4 | 5 |
| $r_{max} = 3 \text{ \AA}$ | | | | | | | | | | | | |
| $e_{abs, mean}$ | 0.0011 | 0.0105 | 0.0160 | 0.0127 | 0.0072 | 0.0045 | 0.0011 | 0.0007 | 0.0008 | 0.0008 | 0.0008 | 0.0008 |
| $e_{abs, max}$ | 0.0033 | 0.0565 | 0.0482 | 0.0436 | 0.0217 | 0.0116 | 0.1122 | 0.0791 | 0.0967 | 0.0855 | 0.0858 | 0.0670 |
| $e_{rel, max}$ | 0.0004 | 0.0060 | 0.0052 | 0.0047 | 0.0023 | 0.0012 | 0.0120 | 0.0085 | 0.0103 | 0.0091 | 0.0092 | 0.0072 |
| $r_{max} = 5 \text{ \AA}$ | | | | | | | | | | | | |
| $e_{abs, mean}$ | 0.0011 | 0.0010 | 0.0027 | 0.0067 | 0.0069 | 0.0045 | 0.0011 | 0.0007 | 0.0008 | 0.0008 | 0.0008 | 0.0008 |
| $e_{abs, max}$ | 0.0033 | 0.0027 | 0.0219 | 0.0220 | 0.0217 | 0.0116 | 0.0433 | 0.0425 | 0.0358 | 0.0428 | 0.0399 | 0.0426 |
| $e_{rel, max}$ | 0.0004 | 0.0003 | 0.0023 | 0.0024 | 0.0023 | 0.0012 | 0.0046 | 0.0045 | 0.0038 | 0.0046 | 0.0043 | 0.0046 |
| $r_{max} = 10 \text{ \AA}$ | | | | | | | | | | | | |
| $e_{abs, mean}$ | 0.0011 | 0.0010 | 0.0010 | 0.0009 | 0.0007 | 0.0010 | 0.0010 | 0.0006 | 0.0007 | 0.0007 | 0.0007 | 0.0007 |
| $e_{abs, max}$ | 0.0033 | 0.0027 | 0.0030 | 0.0028 | 0.0024 | 0.0063 | 0.0120 | 0.0109 | 0.0104 | 0.0104 | 0.0102 | 0.0109 |
| $e_{rel, max}$ | 0.0004 | 0.0003 | 0.0003 | 0.0003 | 0.0003 | 0.0007 | 0.0013 | 0.0012 | 0.0011 | 0.0011 | 0.0011 | 0.0012 |

**Figure 5.**
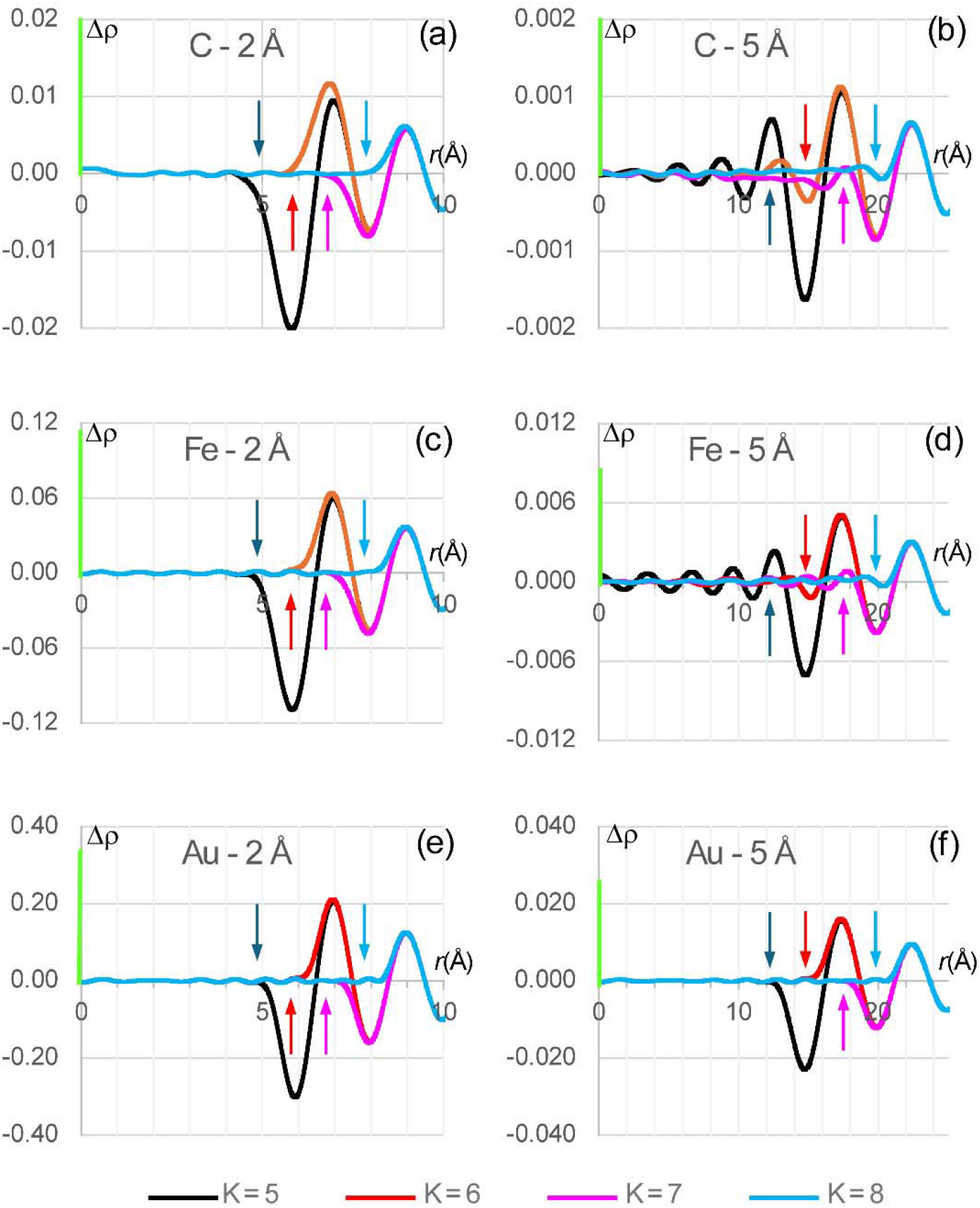
Difference Δ*ρ(r*), in e Å^-3^, as a function of the distance r to the atomic center, between the atomic image (8) – (9) for atoms of C, Fe and Au, and several approximations (1) to it, varying the number of terms *K*. A green bar at the Δ*ρ* axis shows the value *ρ*(0) of the respective atomic image at its center divided by 100. Δ*ρ*(*r*) is computed for images of resolution *d*_*min*_ of 2 Å and 5 Å up to the distance limit 5*d*_*min*_. Colored arrows matching curves’ colors indicate the value of the largest *R*^(*k*)^ for the given approximation. Terms with *R*^(*k*)^ ≤ *r*_*max*_ (considered here as 2.5*d*_*min*_, 3.0*d*_*min*_, 3.5*d*_*min*_, and 4.0*d*_*min*_) may be insufficient to produce an accurate approximation for the points close to *r*_*max*_ and one or two next terms beyond *R*^(*k*)^ may notably reduce the error.

After *R*^(6)^ is included, this error is reduced at the interval [0; 5Å] while still be increased approximately at approaching to 6 Å (*q*_*max*_ = 3.0). Similarly, to improve this approximation up to this limit, an extra term beyond 6 Å, *R*^(7)^ = 6.97, is required (Fig. 5a, curve in magenta).

Depending on atom type and resolution, a need for such extra terms may be weaker (Figs. 5c, e, f) or stronger (Figs. 5b, d). For example, for the image of a carbon atom at the resolution *d*_*min*_ = 5 Å (Fig. 5b), compare the residual error up to the distance 12.5 Å (*q* = 2.5)and 15.0 Å (*q* = 3.0) for the series with K = 5, 6 and 7 terms with *R*^(5)^ 12.13, *R*^(6)^ 14.84, and *R*^(7)^ 17.41, respectively.

The extended interval is defined as *r*_*extended*_ = (*q*_*max*_ + *q*_*add*_)*d*_*min*_ with an extra parameter *q*_*add*_ which, as *q*_*max*_, is an integer of a half-integer. Usually, adding a single term with 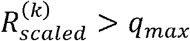 (or, respectively, *R*^(*k*)^ >*r*_*max*_) is sufficient. In most of cases, this term belongs to the interval (*q*_*max*_,*q*_*max*_ + 0.5). Occasionally a larger interval (*q*_*max*_,*q*_*max*_ + 1.0) may be required, which also may allow to identify not one but two following terms, depending on details of a particular curve.

To model all ripples in the scaled image up to the distance *q*_*max*_ by (1), we define the maximal allowed starting approximation error 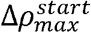 to be equal to 90% of the absolute value of the last ripple at the scaled interval [0;*q*_*max*_]. This guaranties that all local extrema of the curve are interpreted by the terms in (1). Refinement of the coefficient value usually reduces this starting approximation error to the value of order of 2·10^-4^ of the maximal value of the atomic image. When *q*_*max*_ approaches to 15, the starting accuracy defined as above falls below 2·10^-4^ and cannot always be assured with a single term per ripple. However, a search for approximation to atomic images and their analysis at such conditions is out of practical interest.

### 2.5. Variable Resolution Map Calculations for Atomic Models

Calculating a Variable Resolution Map for an atomic model requires assigning a resolution to each atom, which can be the same for all atoms. Next, for each atom type and its associated resolution, the approximation coefficients are retrieved from pre-calculated tables as described in 2.4.2. The map sampling procedure is then applied, consisting of a loop over all atoms to add each atom’s contribution to the map grid nodes that fall within a sphere of a given radius *r*_*max*_ around atomic center. The radius of this sphere depends on the resolution, the atomic displacement parameter, and the purpose of the map being calculated. Choosing a specific radius value involves a trade-off between map accuracy and computation time. A larger radius increases the map’s accuracy; however, it also requires adding atomic contributions to more grid points, resulting in slower calculations. Possible choices of radii and its impact on map accuracy is discussed in Section 3. The local resolution, that needs to be defined for each atom, can be obtained using *phenix*.*local_resolution* tool, which computes the resolution at each map voxel and can optionally transfer that information to each atom in the supplied atomic model using the extended mmCIF format field *_atom_site*.*phenix_resolution*, now supported by CCTBX and Phenix.

## 3. Results

### 3.1. Tabulating image approximation parameters for X-ray and electron diffraction

As mentioned in section 2.4.2, in highly repetitive calculations such as atomic model refinement, calculating image approximation parameters can become a computational bottleneck. Therefore, it is desirable to pre-calculate and store these values, allowing for easy and fast lookup rather than recomputing them on the fly. To this end, we calculated the *A*^(*k*)^, *B*^(*k*)^, and *R*^(*k*)^ values for all entries (atoms and ions) in the X-ray (Waasmaier & Kirfel, 1995) and electron scattering function (Peng *et al*., 1996; Peng, 1999) tables available in CCTBX across a resolution range from 1 to 10 Å, with the step size rationalized in section 2.4.2 within the interval [0; 10 Å].

These tables contain coefficients allowing to calculate approximations up to the distance *r*_*max*_ = 5*d*_*min*_. All the respective approximations contain one term per ripple (10 terms in total) and assure the maximal difference between this approximation and the atomic image (8-9) is of order of 2· 10^-4^ of the values of the atomic image in its center. An extended version of the tables up to the limit *r*_*max*_ = 10*d*_*min*_ (22 terms per approximation) has been also calculated.

### 3.2. Test aims, data, metrics and expected sources of errors

In what follows, we describe numerical tests that aim to address two main questions pertinent to the practical use of the described methodology, namely: 1) the accuracy of the Variable Resolution Map (*ρ*_*VRM*_) and 2) the time needed to compute it, both questions relative to the standard procedure of computing a finite resolution map using Fast Fourier Transform (FFT) with structure factors calculated using the direct summation algorithm (referred to as *reference, ρ*_*FT*_). All maps were generated on a regular (128, 144, 160) grid. Integrals (8-9) were calculated numerically using the standard Simpson’s rule formula computed on a sufficiently fine regular grid on the inverse resolution (2000 equal intervals for 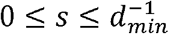).

For the tests with artificially constructed atomic models, including a two-atom model and a protein (PDB code 1crn), we placed these models into an orthogonal large unit cell of size 64×72×80 Å with P1 symmetry. Each of these models aimed at depicting particular source of errors detailed below. We used the 6-Gaussian approximations by Waasmaier & Kirfel (1995) for X-ray diffraction scattering functions, as available in CCTBX.

Three principal sources of discrepancies between *ρ*_*FT*_ and *ρ*_*VRM*_ are:

1. Errors in the map *ρ*_*VRM*_ due to using approximation (1) as opposed to the similar map *ρ*_*image*_ calculated with exact atomic images (8-9);
2. Including contributions within finite distances *r*_*max*_ from atoms only as opposed to computing atomic contribution to all grid nodes;
3. Contributions arising from adjacent by periodicity cells; these contributions are inherently present in the *ρ*_*FT*_ and crystallographic unit cells but are absent in atomic images computed in non-periodic volumes, *ρ*_*image*_, as in cryoEM.

To evaluate discrepancies between the reference and *ρ*_*image*_ or *ρ*_*VRM*_ maps, we used the following metrics:

- Mean and maximal values of the absolute error, defined as the absolute difference computed point by point between the maps for all selected grid nodes, denoted as e_abs,mean_ and e_abs,max_.
- Mean and maximal values of the relative error, defined as the absolute difference computed point by point between the maps for all selected grid nodes, divided by the value 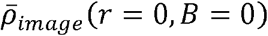 of the heaviest atom in the model, denoted as e_rel,mean_ and e_rel,max_.

These errors were evaluated separately inside and outside the molecular, defined as an accumulation of grid nodes within 2.5 Å radius from the atom positions.

### 3.3. Effect of the image truncation distance

#### 3.3.1. Two-atoms model

In this test, we placed two carbon atoms with a zero B-factor into the unit cell. The first atom was positioned at the center of the unit cell, while each of the three coordinates of the second atom was shifted by *m* Å, where *m* = 0, 1, …, 5 Å. This resulted in six two-atom models, with the atoms spaced apart by 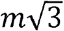 Å and located exactly on the grid nodes. The molecular mask was calculated with *r*_*atom*_ 2.5 Å. Three versions of *ρ*_*image*_ (8-9) were calculated with atomic contributions truncated at *r*_*max*_ values of 3, 5 and 10 Å. Figure 6 schematically illustrates some of these cases.

**Figure 6.**
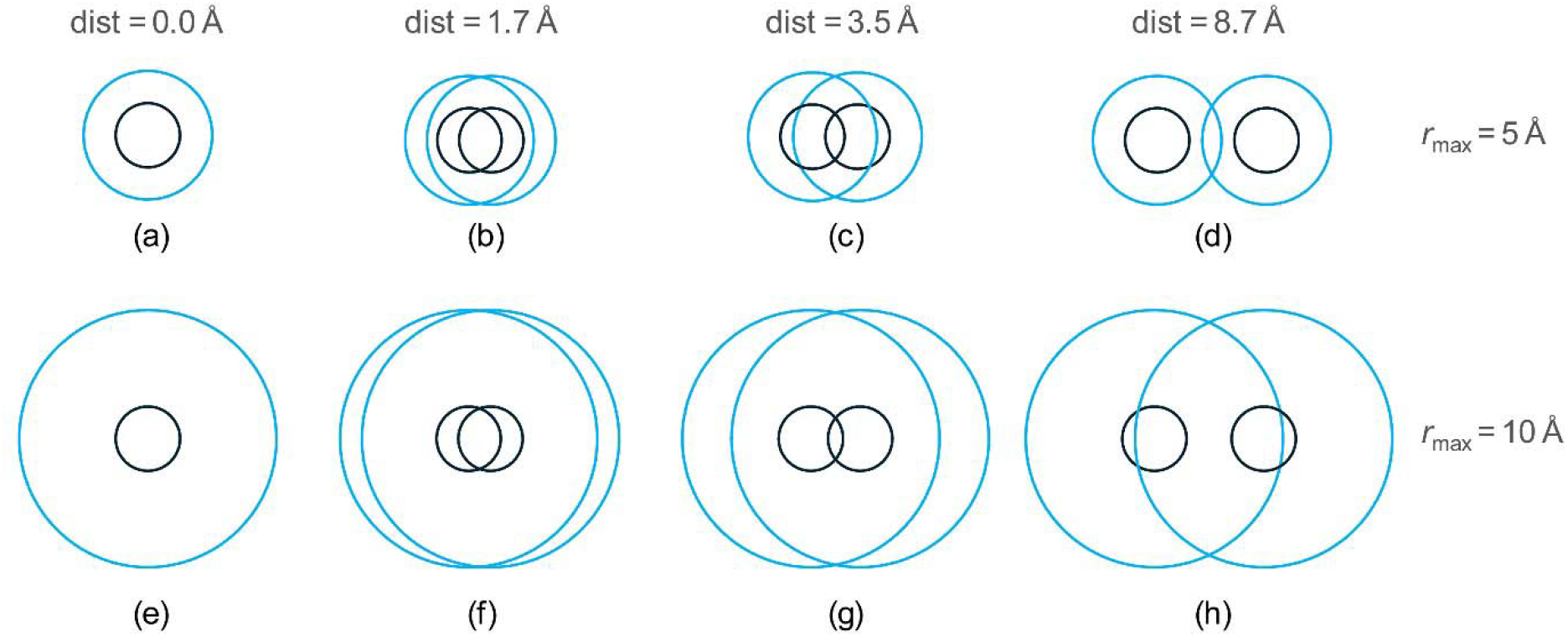
Schematic illustration of different atomic arrangements in some of the two-atom models. Black circles indicate atomic regions (masking radius *r*_*atom*_ = 2.5 Å). Blue circles with the radius *r*_*max*_ indicate regions where atoms contribute to.

For the maps calculated with the exact images (8-9), when the two atoms coincide (Figs. 6a, 6e), the discrepancies between the two maps within the molecular region are attributed solely to contributions from neighboring cells, which the*ρ*_*FT*_ map includes but *ρ*_image_ and *ρ*_*VRM*_ do not. These discrepancies, independent of *r*_*max*_, are relatively small, measuring 0.0011, 0.0033, 0.0004 for e_abs,mean_ and e_abs,max_ and e_rel,max_, respectively. The error outside the molecular region is the sum of the former one and that of the image cut-off, with the latter one dominating. Obviously, for all cases with *r*_*max*_ > *r*_*atom*_, the error outside molecular region decreases with *r*_*max*_ (Table 1, right part, column ‘0’).

When the distance between atomic centers is relatively small compared to *r*_*max*_ (e.g., the distance *dist* = 1.73 Å and *r*_*max*_ equal to 5 and 10 Å), atomic images contribute to all points inside the molecular region (Figs. 6b, f) maintaining the same map difference (Table 1, columns ‘1’). For an intermediate interatomic distance values, one atom can contribute or not to the maps for the second atom (Table 1, columns ‘2’, ‘3, ‘4’) depending on the *r*_*max*_ value (comparer Fig 6c with 6g). For a large interatomic distance, for example 8.7 Å (Figs. 6d, h), the error inside the molecular mask increases (Table 1, columns ‘5’), but less for larger values *r*_*max*_ since the contributing atomic image 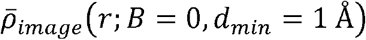 (8) is cut at lower value. These numbers show that the contribution of atoms from neighboring cells can be ignored if the molecule is well isolated in the cell (placed in sufficiently large box), as this is usually the case for cryo-EM studies.

Another important observation is that substitution of *ρ*_*image*_ with *ρ*_*VRM*_ leads to a rather small difference with the maximal deviation of the order of 0.05% with respect to the value of the atomic image in its origin (data not shown). The main conclusion of these tests is that the principal difference between *ρ*_*VRM*_ and *ρ*_*FT*_ is due to the truncation of the atomic images.

#### 3.3.2. Parameters for VRM maps calculation

As pointed out in §2.4.3 and demonstrated in §3.3.1, the accuracy of *ρ*_*VRM*_ depends on *q*_*max*_ and *q*_*add*_ parameters, the choice of which, in turn, depends on the resolution and atomic *B* factors, as these two blur atomic images. Statistics over the structures in the Protein Data Bank (Berman *et al*., 2000; Burley *et al*., 2025) shows the dependency of structure-mean *B* factor as function of resolution as depicted in figure 7. We used this as a guide to sample *q*_*max*_ and *q*_*add*_ for pairs of (resolution *d*_*min*_, *B*) that follow this curve.

**Figure 7.**
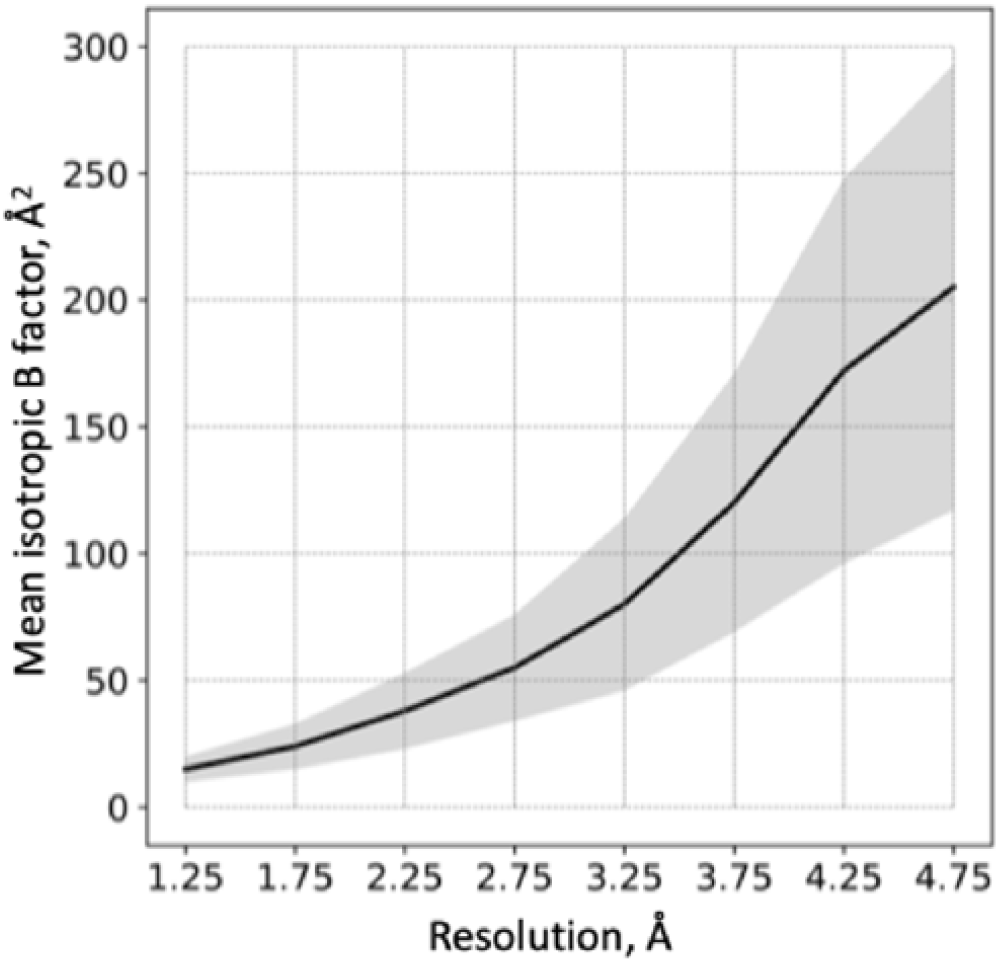
Structure-mean B factor as function of resolution for models in PDB (black curve). Shaded area shows the standard deviation. Marks on x-axis show mid-points of resolution bins in which averaging of B factors was done: (1,1.5), (1.5,2),…, (4.5, 5) Å.

We observed that expectedly, using *q*_*add*_ = 1.0 instead of *q*_*add*_ = 0.5 systematically improved the values of the statistical characteristics but rather marginally, and further increasing to *q*_*add*_ = 1.5 did not change them anymore (data not shown).

We observed also that in all cases *q*_*max*_ = 1.0 and *q*_*add*_ = 0.5 are sufficient to achieve cross-correlation *CC*_*mask*_ (Urzhumtsev *et al*., 2014) between *ρ*_*FT*_ and *ρ*_*VRM*_ greater than 0.99 for a crambin model (Teeter, 1984; PDB code 1crn) placed in the center of unit cell box as defined in §3.2. There is no objective reason not to generalize this finding to any atomic model that follow Figure 7. The maximum local map error is illustrated by Figure 8a.

**Figure 8.**
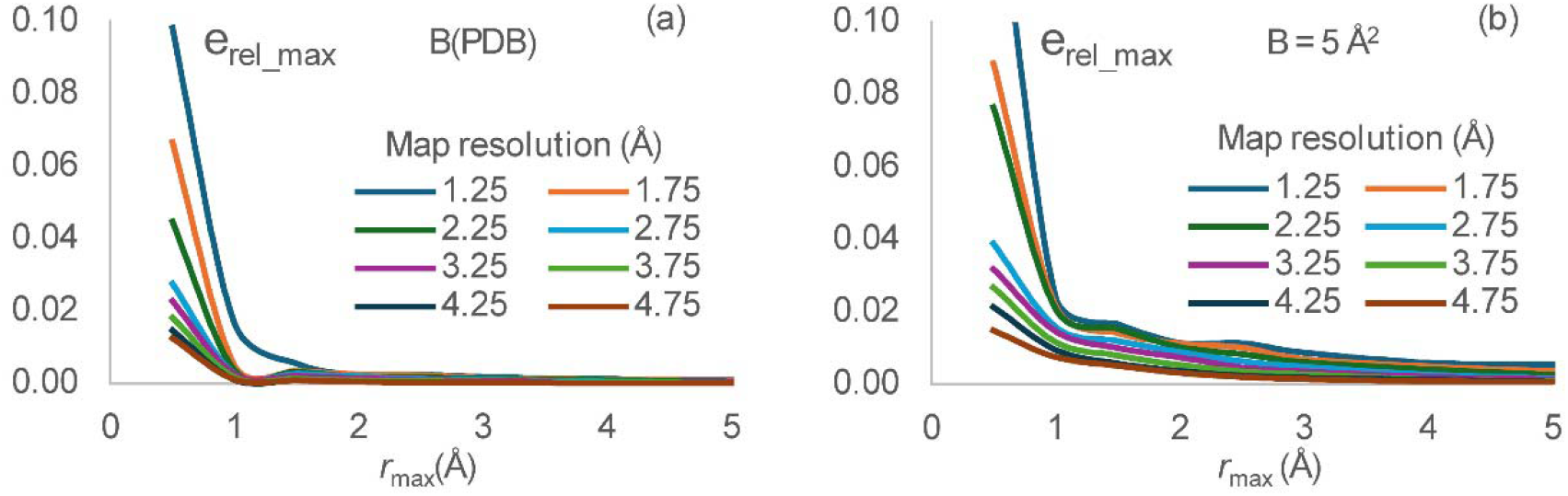
(a) Relative error e_rel,max_ between *ρ*_*FT*_ and *ρ*_*VRM*_ maps inside the molecular mask for the protein model, shown as a function of the image *r*_*max*_; the maps are calculated for different map resolution *d*_*min*_ with *q*_*αdd*_ = 0.5; the *B* values are assigned according to main values from Figure 7. (b) The same as (a) but with all *B* = 5 Å .

Using sharpened maps is common in cryoEM, in which case the mean refined *B* value may not follow the curve in Figure 7, rather the refined B may be well below 10 and typically close to zero; in this case *q*_*max*_ may need to be increased to 1.5-2 to maintain correlation match between *ρ*_*FT*_ and *ρ*_*VRM*_ greater than 0.99; see Figure 8b for the respective maximum local map error.

### 3.4. Runtime illustrations

The CPU time increases cubically with the distance *r*_*max*_, and inversely cubically with the map calculation grid step. The choice of grid step depends on the specific task. A grid step smaller than *d*_*min*_/2 does not provide new information (Nyquist, 1928; Shannon, 1949).

In the following test, we compare the times needed to compute *ρ*_*FT*_ and *ρ*_*VRM*_ maps, computed on grids with step of *d*_*min*_/2 for a series of models with atom counts ranging from 500 to 50,000. The *ρ*_*FT*_ was computed using Fast Fourier Transform (FFT), with structure factors also calculated using FFT and all default parameters of CCTBX, at resolutions of 2 Å (most common for X-ray crystallography) and 4 Å (cryo-EM), utilizing *q*_*max*_ and *q*_*add*_ defined in §3.3.2. All models were placed in a P1 unit cell with dimensions such that the buffer between any atom of the model and the nearest adjacent cell is no less than 5 Å. The specific identity of these models isn’t of particular interest in the context of this test.

Figure 9 shows the time ratios *ρ*_*FT*_ : *ρ*_*VRM*_ for these scenarios as a function of the number of atoms at two different resolutions. For a given resolution, the ratio increases slightly with the number of atoms (approximately logarithmically). The proposed method, applied at a reasonably high resolution of about 5 Å, not only accommodates variations in local resolution but also provides a gain in CPU time. Notably, there is a similarity between the curves calculated at different resolutions. However, at lower resolutions, there is no additional gain in CPU time (nor any significant loss).

**Figure 9.**
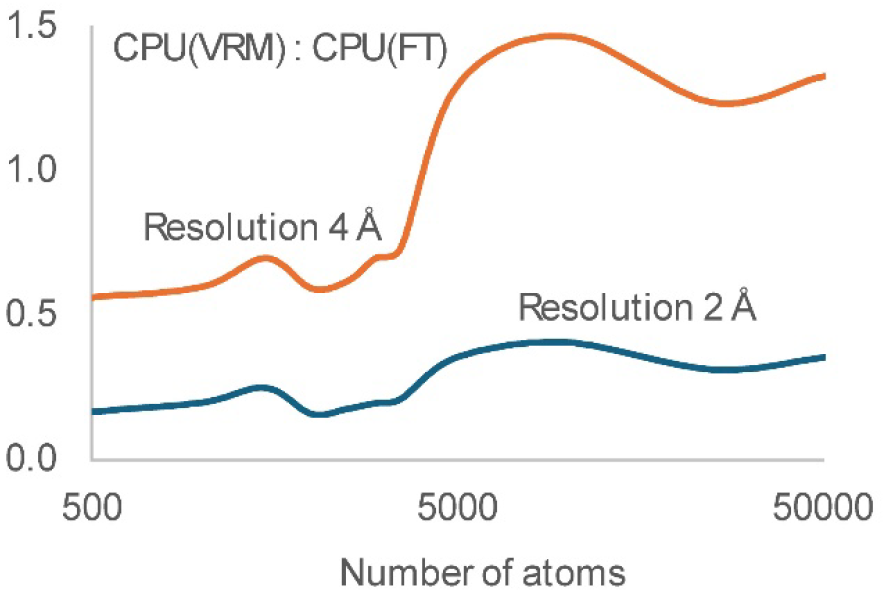
Ratio of the CPU time to calculate a map by VRM (this work) to that for a map calculation by the traditional 3-steps FT-based procedure. The ratio is shown as a function of the number of atoms for the maps calculated at a resolution of 2 and 4 Å with the default parameters as described in the text.

### 3.5. VRM in action: model-versus-map validation in cryoEM

One of the impactful applications of VRM maps mentioned in the Introduction is the validation of atomic models against cryoEM maps. The most accurate and widely used metric for this purpose is CC_mask_, which is the cross-correlation coefficient between the model-calculated map and the experimental map. The strength of this metric lies in its ability to consider all atomic model parameters, which is critical for accurately representing the model in the computed map, and to calculate this map at a resolution that matches the overall consensus resolution of the experimental map. A major drawback of this metric is that it assumes uniform resolution across the entire map, as the map is computed using the three-step procedure outlined in the Introduction. The introduction of VRM maps eliminates this drawback because the resolution of a VRM map can vary at the atomic level.

To illustrate this, we selected an atomic model of the serotonin-bound 5-HT3A receptor in Salipro and its corresponding cryoEM maps (PDB code: 6Y5A, EMDB code: 10692). The resolution of this map, as calculated using *phenix*.*local_resolution* with the supplied half-maps, varies between 2.6 and 6.0 Å, with an average resolution of 3 Å (Fig. 10a). We then calculated the overall value of CC_mask_ for the entire model and its breakdown per individual residue, comparing the experimental map with each of the two: *ρ*_*FT*_ and *ρ*_*VRM*_.

**Figure 10.**
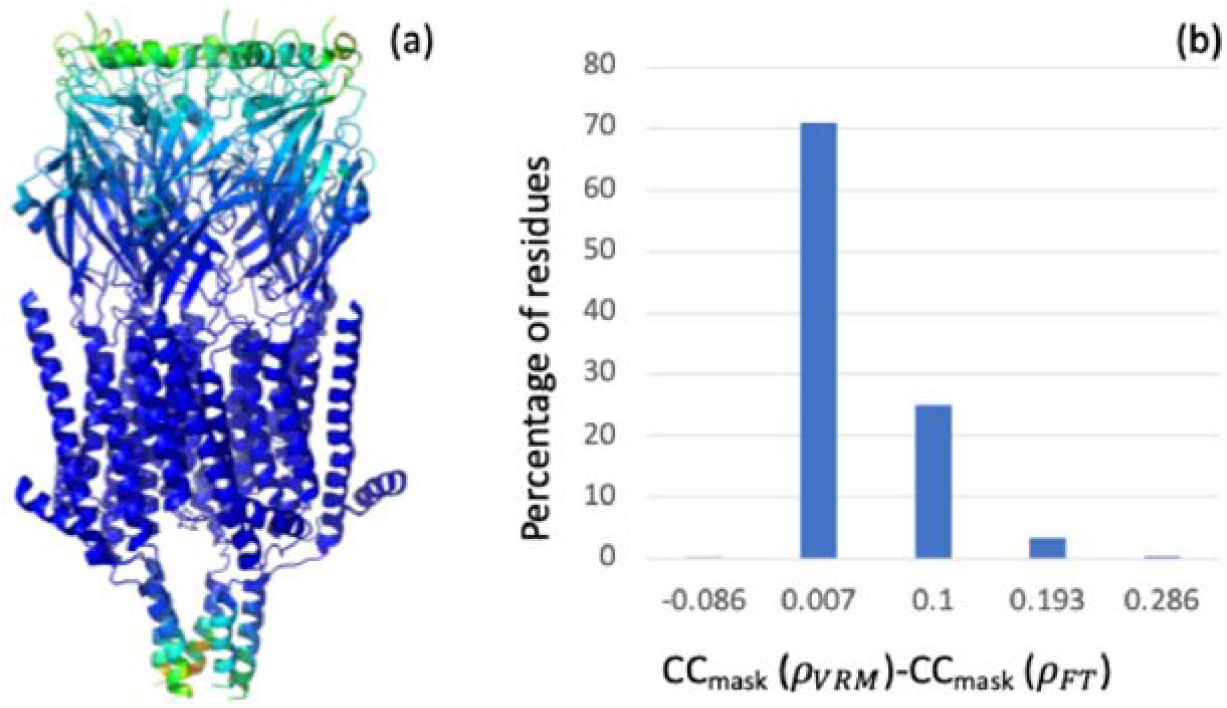
(a) 6Y5A model colored by resolution, ranging from 2.6 Å (blue) to 6 Å (orange). (b) Histogram of differences between CC_mask_ values computed per residue using *ρ*_*FT*_ and *ρ*_*VRM*_ maps.

For the calculation of the *ρ*_*FT*_ map, the resolution for each atom was assigned using the resolution map computed by *phenix*.*local_resolution*, employing tri-cubic interpolation of the map values from surrounding grid nodes of the resolution map onto atomic positions. Figure 10b shows the histogram of differences between CC_mask_ values calculated using *ρ*_*FT*_ and *ρ*_*VRM*_ per residue. We note the improvement of the overall CC_mask_ computed with *ρ*_*VRM*_ (0.6069) over the value computed with *ρ*_*FT*_ (0.5883).

## 4. Discussion

In this manuscript, we describe the implementation of methods in CCTBX and Phenix that enable the calculation of finite-resolution maps, where the resolution can vary from atom to atom within the molecular volume. These methods are based on analytical expressions originally developed by Urzhumtsev & Lunin (2022). This implementation also includes the gradients of model-calculated maps with respect to atomic model parameters, enabling their use in atomic model refinement.

This work led to the addition of resolution as an atomic property in mmCIF files (specifically, the *_atom_site*.*phenix_resolution* field) generated by Phenix. This simplifies the dissemination and exchange of this information between Phenix programs where it is needed. We have tabulated the coefficients for all elements known to X-ray and electron scattering factor tables for faster, non-redundant calculations.

Numerous self-consistent tests presented here demonstrate that VRM maps accurately match those traditionally computed using FFT, which served as the reference when the resolution is distributed uniformly. Additionally, our efforts to improve the convergence of the refinement of image approximation parameters yielded a significant finding: it is possible to express atomic images in such a way that they appear nearly identical for all atoms at all resolutions.

All code and Python scripts used to generate the test results are available in the maptbx.vrm module of the CCTBX library.

## Acknowledgments

PDA and PVA acknowledge funding from the National Institutes of Health (R24GM141254), as well as support from the Phenix Industrial Consortium and the US Department of Energy under Contract No. DE-AC02-05CH11231. AU acknowledges Instruct-ERIC and the French Infrastructure for Integrated Structural Biology FRISBI (ANR-10-INBS-05) and the support and the use of resources of Instruct-ERIC through the R&D pilot scheme APPID 2683. This work would not have been possible without the close collaboration and numerous fruitful discussions between one of the authors (A.G.U.) and Vladimir Y. Lunin, whose insights were invaluable. We thank Oleg Sobolev (MBIB, LBNL) for extending mmCIF support to define peratom resolution.

